# Melatonin alters fluid phase co-existence in POPC/DPPC/cholesterol membranes

**DOI:** 10.1101/2020.04.03.015966

**Authors:** Nanqin Mei, Morgan Robinson, James H. Davis, Zoya Leonenko

## Abstract

The structure and biophysical properties of lipid biomembranes are important for normal function of plasma and organelle membranes, which is essential for proper functioning of living cells. In Alzheimer’s disease (AD) the structure of neuronal membranes becomes compromised by the toxic effect of amyloid-*β* (A*β*) protein which accumulates at neuron synapses, resulting in membrane perforation and dysfunction, oxidative stress and cell death. Melatonin is an important pineal gland hormone that has been shown to be protective against A*β* toxicity in cellular and animal studies, but the molecular mechanism of this protection is not well understood. It has been shown that melatonin can interact with model lipid membranes and alter the membrane biophysical properties, such as membrane molecular order and dynamics. This effect of melatonin has been previously studied in simple model bilayers with one or two lipid components, we consider a more complex ternary lipid mixture as our membrane model. In this study, we used ^2^H-NMR to investigate the effect of melatonin on lipid phase behaviour of a three-component model lipid membranes composed of 1,2-dipalmitoyl-*sn*-glycero-3-phosphocholine (DPPC), 1-palmitoyl-2-oleoyl-*sn*-glycero-3-phosphocholine (POPC) and cholesterol. We used deuterium labelled palmitoyl-*d*_31_ in POPC-*d*_31_ and DPPC-*d*_62_ separately, to probe the changes in hydrocarbon chain order as a function of temperature and varying concentrations of melatonin. We found that melatonin concentration influences phase separation in these ternary mixtures somewhat differently depending on whether POPC-*d*_31_ or DPPC-*d*_62_was used. At 5 mol% melatonin we observed phase separation in samples with POPC-*d*_31_, but not with DPPC-*d*_62_. However, at 10 mol% melatonin phase separation was observed in both samples with either POPC-*d*_31_ or DPPC-*d*_62_. These results indicate that melatonin can have a strong effect on membrane structure and physical properties, which may provide some clues to understanding how melatonin protects against A*β*.

**SIGNIFICANCE:** Melatonin has been shown to be protective against A*β* pathology in animal and cellular studies. Although the mechanism of this protection is not well-understood, melatonin’s membrane-active properties may be important in this regard. In this work solid-state deuterium nuclear magnetic resonance was used to study the effect of melatonin on the POPC/DPPC/cholesterol model membranes. Specifically, we showed that melatonin modifies lipid hydrocarbon chain order to promote phase separation. This knowledge helps to explain the role of melatonin in lipid domain formation and may provide a deeper understanding of the mechanism of melatonin neuroprotection in AD.

## INTRODUCTION

The plasma membrane is an essential cellular structure and its structure and function is sensitive to lipid composition which affects biophysical properties of the lipid bilayer and can modulate membrane protein function. Due to the complexity of natural cellular membranes, which are composed of hundreds of different lipids including phospholipids and cholesterol, simpler model lipid membranes composed of synthetic lipids are widely used to mimic the plasma membrane in order to study membrane properties (1–3). Changes in the properties of the biomembrane have important implications in health and disease. An important example is Alzheimer’s disease (AD), where it has been shown that the brain lipid composition changes with age (4, 5) and that these changes may lead to an increase of amyloid-*β* (A*β*) toxicity and damage to the membrane (2, 6). With age, the accumulation and aggregation of A*β* deposits on the membranes in the brain leads to cell membrane damage, increased membrane permeability, disruption of membrane potential, oxidative stress and ultimately cell death (7, 8). Thus, it is important to understand the relationship between membrane composition, membrane properties and damage to membrane integrity. Specifically to AD, detailed understanding of the role of these membrane changes may help to identify ways to protect the membrane from A*β*-induced damage. Melatonin is a small membrane-active molecule that has been shown to protect against A*β*; however, the molecular mechanism of this protection is not well understood. Understaing the protective role of melatonin in AD may guide new preventive strategies in AD.

Melatonin is a tryptophan-derived pineal gland hormone that is important for regulating circadian rhythm but it is known that its levels decrease with age (9). Moreover, melatonin levels and circadian rhythms have been found to be disrupted in Alzheimer’s disease (AD) patients, as compared to age-matched controls (10, 11). Melatonin has been shown to alleviate these symptoms in some clinical studies (12), though results are mixed (13). In animal studies of AD it has been demonstrated that melatonin is a potent anti-oxidant and is neuroprotective against A*β* toxicity at the cellular level (14–18). Interestingly, the protective effect of melatonin was found to be receptor independent (19), yet the complete molecular mechanisms are not well-understood. The direct incorporation of melatonin into the neuronal membranes may be an important part of these protective mechanisms. Melatonin is a membrane-active molecule and its role on the membrane properties has ben studied in model lipid membranes using a variety of different approaches including X-ray and neutron scattering (1, 20), Fourier transform infrared (FT-IR) spectroscopy (21), Langmuir-Blodgett trough (22), molecular dynamics (MD) simulations (1, 22), and nuclear magnetic resonance (NMR) spectroscopy (21, 23), showing that melatonin has pronounced biophysical effects on simple lipid systems, composed of single or dual lipids,. In this work, we used a ternary lipid membrane model to better mimic the natural cellular membranes and to further understand melatonin’s effects on the membrane structure and properties.

Unevenly distributed molecular constituents produce heterogenous surface topology in cellular membranes; the resulting variable domains are likely an essential factor that impacts membrane function (24). Phase separation refers to the behaviour of heterogeneous lipid mixtures that contain both highly ordered domains, enriched in cholesterol and containing predominantly saturated long acyl chain lipids, and more fluid domains with a higher proportion of unsaturated lipids. These different membrane domains have been proposed to compartmentalize different cellular processes, as suggested by the membrane raft hypothesis (25). The coexistence of liquid-ordered (*𝓁*_*o*_) and liquid-disordered (*𝓁*_*d*_) phases and lipid domains has been observed extensively in model lipid systems using a variety of biophysical techniques (for example, nuclear magnetic resonance, neutron scattering, fluorescence microscopy, and atomic force microscopy) and there is increasing evidence of nanodomains in living systems (1, 26–28). In general, domains rich in cholesterol and long-chain saturated lipids have a higher degree of molecular order compared to the whole membrane system (3). Cholesterol has a particularly interesting bidirectional effect on lipid membranes depending on its amount: at sufficiently high concentration it preserves the *𝓁*_*o*_ phase at high temperature while suppressing the *𝓁*_*o*_ to solid phase transition at low temperature (3, 29, 30).

Solid state ^2^H NMR has been widely employed to study these phase equilibria phenomena in lipid membrane systems (3, 30, 31). In this work we use ^2^H NMR to study the influence of melatonin on the phase equilibria of complex mixtures containing POPC, DPPC and cholesterol. By substituting POPC or DPPC with corresponding deuterated lipids POPC-*d*_31_ or DPPC-*d*_62_, we show that phase separation induced by melatonin at low concentration is apparent when POPC is labelled but not DPPC, suggesting a stronger interaction of melatonin with the more disordered lipid membrane regions containing higher abundance of POPC. This result may also demonstrate an effect caused by the choice of chain perdeuteration which highlights the importance of isotopic labelling in lipid membrane studies. Overall, the influence of melatonin on the phase equilibrium phenomena in this tertiary model lipid system suggests it may induce phase separation and that melatonin has a preference for more intrinsically disordered lipids that have a larger area per molecule within the membrane.

## MATERIALS AND METHODS

### Sample Preparation

All lipids were purchased from Sigma-Aldrich and used without further purification. Lipid stock solutions were prepared using chloroform at a concentration of 10 mg/ml. A lipid model system composed of POPC, DPPC, and cholesterol at a molar ratio of 3:3:2 was used throughout. Two types of deuterium labelled lipid mixtures, POPC-*d*_31_/DPPC/cholesterol and POPC/DPPC-*d*_62_/cholesterol, were prepared. Melatonin was added to each type of lipid mixture at a molar ratios of 0, 5 and 10% to obtain a sequence of samples. Also, pure POPC-*d*_31_ and pure DPPC-*d*_62_ samples were tested to determine the effect of perdeuteration to the lipid melting temperature.

To prepare the lipid samples, standard solutions were mixed in a round bottom flask, according to the ratio desired while insuring the deuterated lipid in each mixture was at least 5 mg. Because the quadrupolar splittings of ^2^H NMR are very sensitive to the concentration of cholesterol in the liquid-crystalline phase, a heterogeneous distribution of cholesterol will lead to a broad distribution of splittings (30). Ethanol has been shown to reduce heterogeneity of lipid mixtures when compared with chloroform (3, 30). Thus, after lyophilization from chloroform overnight, ethanol was used to redissolve the lipid mixtures. The sample was then freeze-dried overnight from ethanol and scraped out from the bottom of the flask. Each sample was transferred into a snap-cap centrifuge tube and resuspended in 50 mM phosphate buffer (pH 7.0), at weight ratio of lipid to buffer of 4:3. The sample was mixed with a thin glass rod and the whole mixture was centrifuged into the bottom of the tube. This process was repeated 2-3 time to create a uniformly-distributed multi-lamellar mixture (30). The whole mixture was transferred into a 3 mm diameter NMR tube by centrifugation through a small hole that was cut on the bottom of the snap-cap tube. The NMR sample tube was sealed with silicone and was given at least 2 hours for the silicone cap to dry out. The entire sample tube was then weighed before and after each use to monitor any possible loss of water during the experiment.

### ^2^H NMR

At the beginning of each series of NMR measurements the sample was heated to an initial set point of 324 K in order to assess the sample homogeneity from the ^2^H NMR spectrum itself. Spectra were then taken as a function of temperature, cooling the set point in 1 K steps from 324 down to 268 K. At each cooling step the temperature was allowed to come to equilibrium at the new set point temperature after which the sample was allowed an additional 10 minutes to equilibrate before the spectrum was taken. Each spectrum consisted of the accumulation of 4096 scans obtained using the quadrupolar echo sequence (32). At the end of each temperature series the samples were then reheated to a set point of 300 K and subsequently to 324 K where spectra were again taken to verify the reproducibility of the spectra and the stability of the sample. Since the sample temperature is not alway exactly equal to the set point temperature, the NMR probe temperature was calibrated using the temperature dependent frequency shift of Pb(NO_3_)_2_ (33, 34). The melting points of DPPC/water and DMPC/water were used as references for the calibration. The temperatures shown in the figure captions and on the figures are the corrected sample temperatures. After all NMR experiments were completed on a given sample the sample was weighed for comparison with the initial weight. In this fashion we were able to insure that no water was lost during any of the measurements.

All NMR experiments were performed on a Bruker Avance II Plus 500 MHz spectrometer operating at a ^2^H NMR resonance frequency of 76.72 MHz. The quadrupolar echo sequence consists of two 90^*o*^ pulses, each 2.9 µs long, separated by an echo forming delay τ of 25 *µ*s. In order to refocus the precession due to the quadrupolar interaction during τ the second pulse is phase shifted relative to the first by 90^*o*^ (32). The relaxation delay between scans was 0.5 s.

Because ^2^H is a spin 1 nucleus, the ^2^H NMR spectrum is dominated by the quadrupolar interaction between the electric quadrupolar moment of the nucleus, *eQ*, and the electric field gradient at the nucleus, *eq* (27). This interaction results in two peaks to the spectrum of a single ^2^H, which are separated by the quadrupolar splitting. The value of the quadrupolar splitting depends on the orientation of the principal axis of the electric field gradient (aligned along the C-^2^H bond in lipids) and the external magnetic field

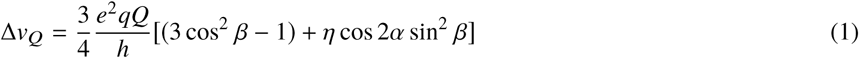

Where the angles α and *β* are Euler angles specifying the orientation of the principal axis system of the electric field gradient tensor relative to the external magnetic field **B**_0_. This expression is significantly simplified for ^2^H in C-^2^H bonds because the asymmetry parameter, *η*, is very small and can be taken to be 0. The maximum quadrupolar splitting for ^2^H occurs when there is no molecular reorientation relative to the magnetic field and is found when the C-^2^H bond is aligned parallel to the field. In that case the observed splitting would be 252 kHz. In fluid lipid bilayers there is a lot of molecular motion so that the quadrupolar splittings observed are greatly reduced. The experimental value of the quadrupolar splitting for a ^2^H nucleus in a C-^2^H bond provides a useful measure of the degree of molecular motion experienced by that segment of the molecule. It can range from a value of 0, observed for rapid isotropic molecular reorientation, to the rigid lattice limit of 252 kHz (27).

Lipid hydrocarbon chain order is much higher for lipids within a liquid ordered, *𝓁*_*o*_, phase domain than for lipids in a liquid disordered, *𝓁*_*d*_, phase domain. For samples made with deuterium labelled lipids (such as DPPC which is deuterium labelled at a single chain position), at temperatures and compositions that are within a two phase, *𝓁*_*o*_:*𝓁*_*d*_, region we would expect to see two very distinct quadrupolar splittings. If the lipid chains are labelled at all positions then, because of the variation of C-^2^H bond order with chain position (27), there will be a superposition of many quadrupolar splittings in the spectra but those lipids which are in the more ordered *𝓁*_*o*_ phase will have much larger quadrupolar splittings than those lipids which are in the relatively disordered *𝓁*_*d*_ phase.

### Quadrupolar Splitting Analysis

The multiple overlapping quadrupolar splittings from the chain methylene groups are frequently difficult to resolve. In order to compare chain order at different temperatures and different sample compositions, we would typically calculate the first moment, *M*_1_, of the ^2^H spectrum since it is proportional to the average quadrupolar splitting (31). However, the chain terminal methyl group splittings are always distinguishable from the methylenes so we also use their quadrupolar splittings in sample comparisons. The methyl group splittings were measured directly as the frequency difference between the peaks in the methyl group powder patterns found near the center of each spectrum.

## RESULTS AND DISCUSSION

The magnitudes of the ^2^H quadrupolar splittings observed in ^2^H NMR spectra of deuterium labelled lipid acyl chains are determined by the rapid motions which partially average the quadrupolar Hamiltonian. The character and time scales of these motions depend on sample temperature and composition. Because of this the ^2^H NMR spectra can be used to study the phase equilibria of lipid mixtures (3, 35). Figure 3a(i) shows the temperature dependence of the ^2^H NMR spectra observed in fully hydrated POPC-*d*_31_ bilayers. These spectra are characteristic of chain perdeuterated phospholipid lamellar dispersions in the ‘fluid’ or liquid disordered phase (*𝓁*_*d*_) typically found in bilayers with little or no cholesterol. For POPC-*d*_31_ the sample remains in the fluid phase to quite low temperatures because the chain melting transition occurs at around -4 *°*C for non-deuterated POPC (36). We find that the transition for this sample of POPC-*d*_31_ occurs at about 263.5 K or approximately -9.5 *°*C.

**Figure 1:**
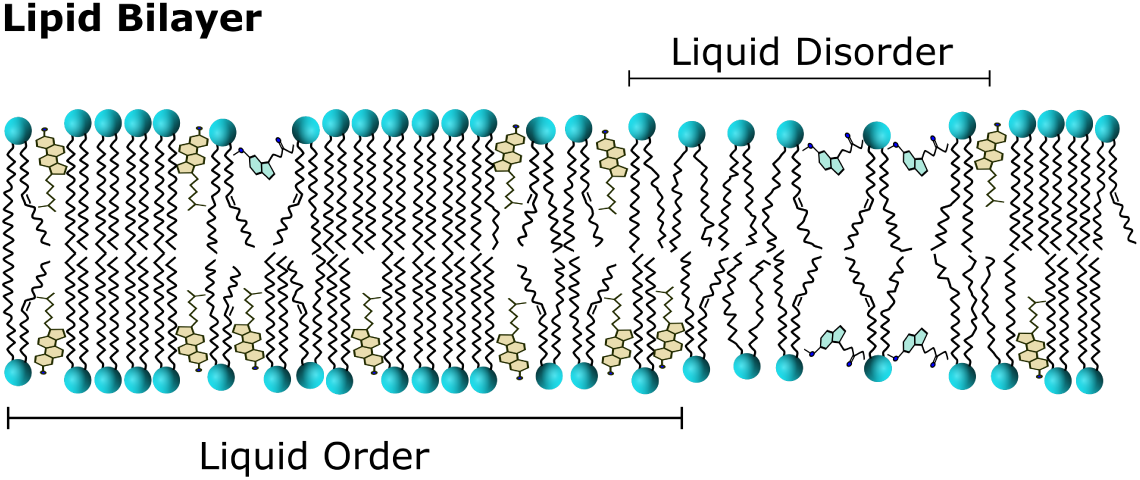
Model system studied in this report: lipid bilayer consisting of POPC, DPPC, cholesterol and melatonin. The coexistance of liquid ordered and liquid disordered phases is represented. Liquid ordered phases have tails that are aligned and more rigid than disordered tails. The orientation of cholesterol and melatonin predicted from MD simulations is shown, with melatonin decreasing the number of lipids per membrane area and cholesterol having opposing effects.

**Figure 2:**
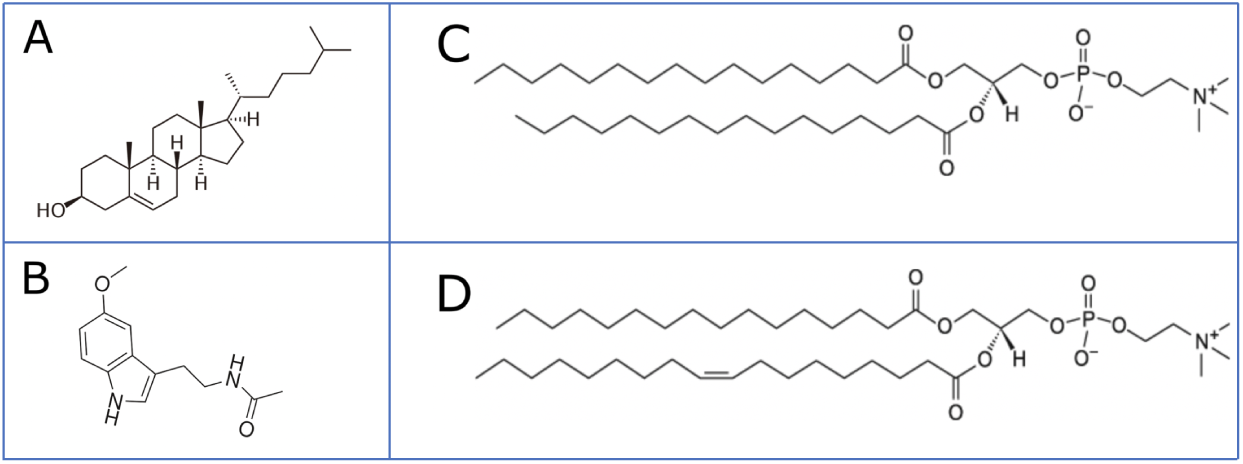
Molecular Structure of A. cholesterol, B. melatonin, C. DPPC and D. POPC

**Figure 3:**
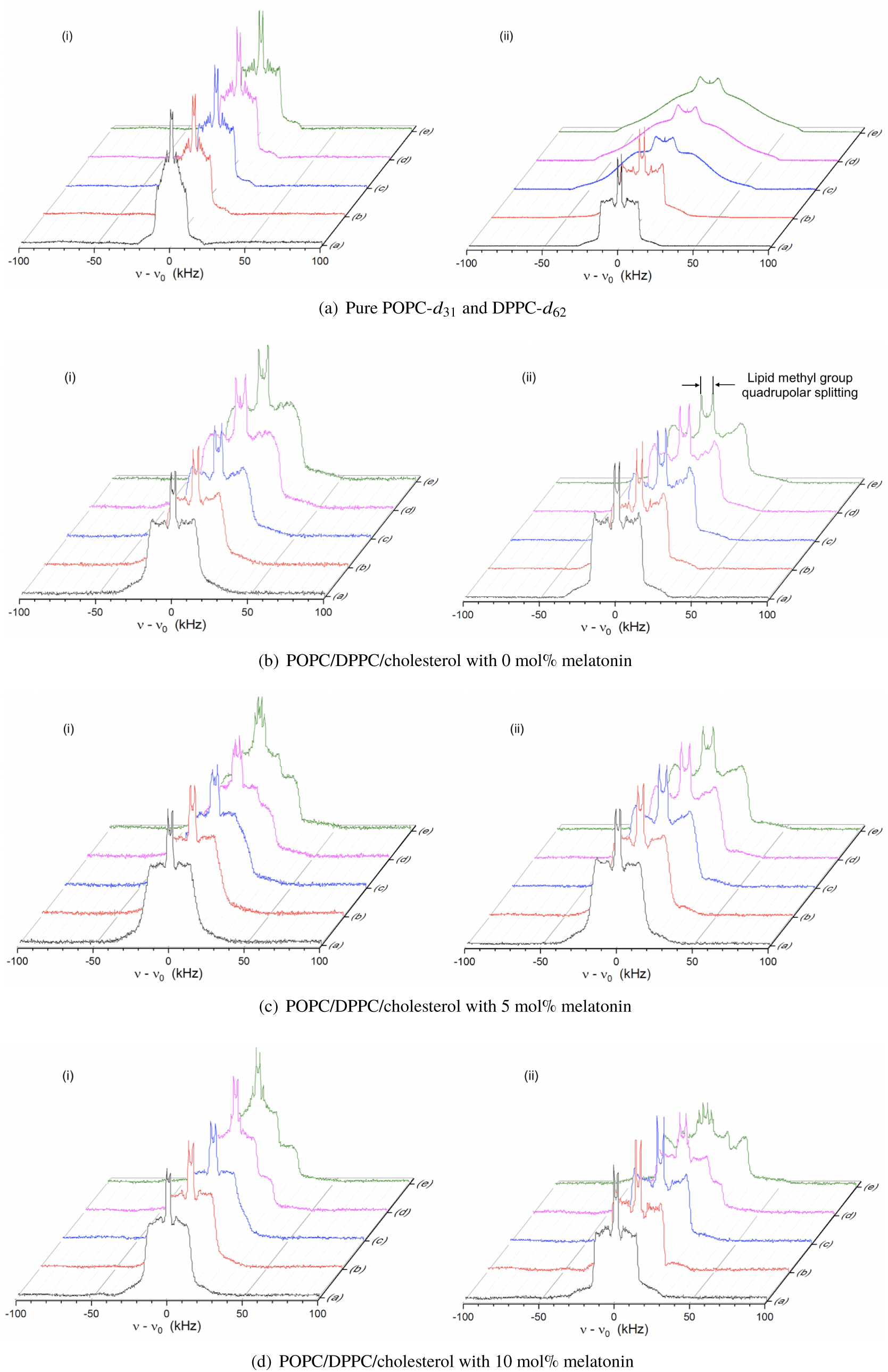
^2^H-NMR spectra of samples with (i) POPC-*d*_31_ or (ii) DPPC-*d*_62_ deuterated lipids. Each individual plot shows the spectra at five different temperatures: (a) 329.2 K, (b) 316.8 K, (c) 303.3 K, (d) 298.4 K, (e) 291.3 K.

Figure 3a(ii) shows the temperature dependence of the spectra of DPPC-*d*_62_. Above about 310 K (37.75 *°*C (30)) the sample is in the fluid phase but below that temperature it enters the gel phase where the ^2^H NMR spectra are much broader due to the suppression of acyl chain *gauche-trans* isomerization resulting from the tighter chain packing in the gel phase (31, 37). The perdeuteration of DPPC lowers its melting temperature by about 4 degrees from 314 K to 310 K (36). Below this chain melting transition the chain motions are greatly restricted and their time scales are considerably longer. Thus, the ^2^H NMR quadrupolar splittings are larger (less motional averaging) and the individual peaks in the spectrum are much broader (due to the slower motions). The spectral line shapes illustrated in Figure 3a(ii) are typical of fully hydrated phospholipid lamellar dispersions in the fluid and gel phases.

In the presence of cholesterol the quadrupolar splittings in the fluid phase increase approximately linearly with increasing cholesterol concentration (38). At sufficiently high cholesterol concentration a phase separation into an *𝓁*_*d*_:*𝓁*_*o*_ two phase region can occur (3, 30). The ^2^H spectra of chain perdeuterated lipids in the liquid ordered *𝓁*_*o*_ phase are similar to those of the *𝓁*_*d*_ phase except that the quadrupolar splittings are much larger (> 50 kHz compared to *≈*22 kHz).

The spectra in Figure 3b(i) are for a sample of POPC-*d*_31_/DPPC/cholesterol with molar proportions of 37.5/37.5/25 while those in Figure 3b(ii) are for POPC/DPPC-*d*_62_/cholesterol with the same molar proportions. For both samples the edges of the spectra have splittings ranging from about 35 to over 55 kHz depending on temperature. These splittings are representative of a fluid lipid bilayer at high cholesterol concentration (38). In a ternary mixture containing a long saturated chain phospholipid, an unsaturated chain phospholipid and a high concentration of cholesterol one might have expected to find *𝓁*_*d*_ : *𝓁*_*o*_ phase coexistence (as for DOPC/DPPC/cholesterol mixtures (3)). Careful inspection of the methyl group part of the spectra in Fig. 3b(ii) reveals the inequivalence of the two hydrocarbon chains for DPPC-*d*_62_ which is a signature of a highly ordered phase (30, 38). Since only one chain is deuterated for POPC-*d*_31_ this feature is absent from the spectra in Fig. 3b(i). However, over the temperature range studied (from -5 to +55 *°*C), there is no evidence of coexistence of macroscopic *𝓁*_*d*_ and *𝓁*_*o*_ phases. This agrees with other reports that do not detect microscale domains of similar POPC/DPPC/cholesterol mixtures (39–42); this is in contrast to ternary systems containing DOPC, with two unsaturated fatty acyl chains, where microscale domains are observed (3, 28). DOPC having two unsaturated hydrocarbon chains, has a lower melting temperature, lower average hydrocarbon chain order parameter, and looser chain packing than POPC (36, 43) which has only a single unsaturated chain. In the presence of high concentrations of cholesterol, these factors seem to be sufficient to lead to macroscopic *𝓁*_*d*_ : *𝓁*_*o*_ phase separation in DOPC/DPPC/cholesterol but not in POPC/DPPC/cholesterol mixtures.

The addition of 5 mol% melatonin (of total lipid, i.e., POPC + DPPC + cholesterol was 95 mol%) to the POPC/DPPC/cholesterol mixtures with the same proportions used for Figure 3c had an interesting effect on the spectra. The sample with POPC-*d*_31_ showed a clear phase separation below about 301.5 *°*C, as shown in Figure 3c(i). On the other hand, the sample with DPPC-*d*_62_, Figure 3c(ii), showed no evidence of phase separation over the temperature range studied. Repeating the experiments with a second set of samples having the same compositions gave identical results. Due to the looser chain packing of unsaturated lipids such as POPC, there is more room at the interfacial region (near the phospholipid headgroup) for melatonin to insert itself than in the more tightly packed regions rich in DPPC and cholesterol. The preferential partitioning of melatonin into regions rich in unsaturated lipids tends to reduce the molecular order even further, enhancing the difference between areas rich in POPC and those rich in DPPC, especially since cholesterol interacts preferentially with saturated chain lipids and dramatically increases their hydrocarbon chain order. This competitive process between cholesterol and melatonin for the two different lipid species is supported by FT-IR and ^1^H-NMR spectroscopy experiments of inverted lecithin micelles containing melatonin and cholesterol that indicate an overlap in the stretching modes of melatonin’s -NH with the -OH of cholesterol and that chemical shifts of the NH protons of melatonin are sensitive to cholesterol content (23). As noted above, chain perdeuteration shifts the phase transition temperature of either POPC-*d*_31_ or DPPC-*d*_62_ downward by about 4 *°*C. This has the effect of increasing the difference between the transition temperatures (and degrees of hydrocabon chain order) of the two lipids in the case of POPC-*d*_31_/DPPC/chol mixtures relative to POPC/DPPC/chol but it decreases those differences for the case of POPC/DPPC-*d*_62_/chol. The simple addition of melatonin at 5 mol% is enough to promote microscale domain formation in one case but not in the other.

At 10 mol% melatonin both samples, whether labelled with POPC-*d*_31_ or DPPC-*d*_62_, show a broad region of *𝓁*_*d*_:*𝓁*_*o*_ two phase coexistence as shown in Figure 3d. As might be expected, these spectra show that at a given temperature within the two phase region there is more of the unsaturated POPC in the *𝓁*_*d*_ phase and more of the saturated DPPC in the *𝓁*_*o*_ phase.

Because of the large number of overlapping ^2^H NMR powder patterns observed when using chain perdeuterated samples it can be difficult to measure and to follow the changes in individual methylene group quadrupolar splittings. However, it is quite easy to measure the splittings of the chain terminal methyl groups as they are considerably smaller than those of the methylenes and their peak intensity is larger.

Figure 4a shows the temperature dependence of the methyl group splittings for samples of pure POPC-*d*_31_ and of pure DPPC-*d*_62_. For POPC-*d*_31_ there is only a gradual monotonic increase in the splitting of the single deuterated methyl group as the temperature is lowered from 332.9 K to 277.8 K. For DPPC-*d*_62_ on the other hand there are a couple of interesting changes. Just below 310 K there is a sharp increase in the splittings of both chain methyl groups as the sample is lowered through the chain melting transition from the fluid into the gel phase. This chain melting transition temperature is about 4 *°*C lower than the non-deuterated DPPC melting temperature at 314 K (36). In the gel phase the two methyl groups are inequivalent, one having a considerably larger splitting than the other. This is because the *<u>sn</u>*-1 chain extends farther into the hydrophobic core than the *<u>sn</u>*-2 chain which is bent where it attaches to the phospholipid glycerol backbone. Even in the gel phase there is significantly more chain mobility at the very center of the bilayer. In addition, at all temperatures the methyl group splitting of POPC-*d*_31_ is smaller than those of DPPC-*d*_62_.

**Figure 4:**
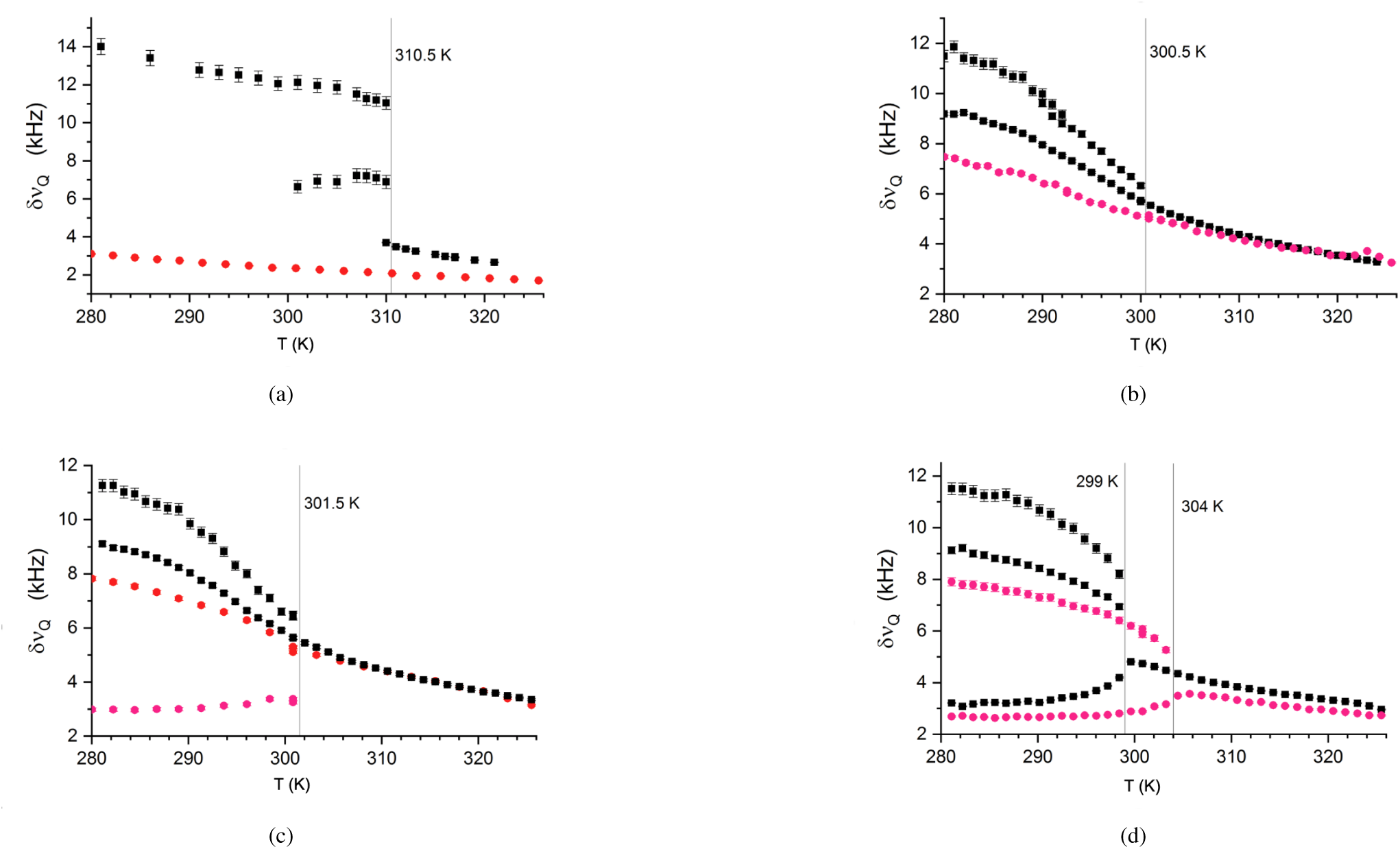
Lipid methyl group quadrupolar splittings of pure POPC-*d*_31_ and DPPC-*d*_62_ (a), and POPC/DPPC/cholesterol with (b) 0 mol%, (c) 5 mol% and (d) 10 mol% melatonin. Black squares represent samples with DPPC-*d*_62_, and red circles represent the samples with POPC-*d*_31_. Lower splittings correlate with less order of the lipid tails, while higher splittings indicate the opposite.

The methyl group splittings for the three component mixtures are shown in Figure 4b. Here, at high temperatures, the POPC-*d*_31_ and DPPC-*d*_62_ methyl splittings are very similar with both gradually increasing as temperature is lowered. At about 300 K we can see the development of the inequivalence of the two DPPC-*d*_62_ chains. This characteristic behaviour is observed whenever the lipid chains are sufficiently highly ordered (38). We do not see any evidence of *𝓁*_*d*_:*𝓁*_*o*_ phase separation, presumably due to the small size of the domains in the absence of melatonin (42).

For samples with 5 mol% melatonin Figure 4c shows that at higher temperatures the POPC-*d*_31_ and DPPC-*d*_62_ methyl splittings are again very similar. For the DPPC-*d*_62_ sample, however, below about 301 K we see the return of the inequivalence of the two DPPC-*d*_62_ methyl groups but see no evidence of two phase coexistence. On the other hand, we see the appearance of two POPC-*d*_31_ methyl group splittings. These are due to the formation of the *𝓁*_*d*_:*𝓁*_*o*_ two phase region observed in the full spectra in Fig. 3c(i). An important signature of *𝓁*_*d*_:*𝓁*_*o*_ two phase coexistence is the fact that in the *𝓁*_*o*_ phase domains the methyl group quadrupolar splitting (in the present case arising only from POPC-*d*_31_) increases as the temperature is lowered due to two factors: the reduced motional averaging at lower temperatures and the increased cholesterol content of the *𝓁*_*o*_ phase as the temperature is lowered. On the other hand, the methyl splitting of the *𝓁*_*d*_ phase decreases as the temperature is lowered due to the decreasing cholesterol content of the *𝓁*_*d*_ phase domains. This behaviour is conclusive evidence for the changing composition of the two phases with temperature.

Finally, in Figure 4d, we show the methyl group splittings for the samples containing 10 mol% melatonin. At high temperatures the methyl splittings for POPC-*d*_31_ are slightly smaller than those of DPPC-*d*_62_ due both to the increased chain disorder of unsaturated lipids relative to saturated lipids and to the higher affinity of melatonin for unsaturated lipids. Comparing the methyl group splittings with 5 mol% melatonin (Figure 4c) to those at 10 mol% melatonin, we also see that the increased melatonin concentration has reduced the hydrocarbon chain order and the methyl splittings.

At and below 304 K, with 10 mol% melatonin, the POPC-*d*_31_ methyl splitting has two components indicating that the phase separation has begun. On the other hand, for the sample with DPPC-*d*_62_ the methyl splittings show that phase separation occurs only for temperatures below about 299 K. Recall, however, that at 5 mol% melatonin no phase separation was observed for the DPPC-*d*_62_ sample over the temperature range studied. As discussed above, chain perdeuteration of POPC-*d*_31_ increases the difference between POPC-*d*_31_ and DPPC chain melting transition temperatures, while chain perdeuteration of DPPC-*d*_62_ reduces the difference between DPPC-*d*_62_ and POPC. Evidently the POPC/DPPC/cholesterol ternary mixture is so close to fluid-fluid phase separation that, in the presence of 5 mol% melatonin, chain perdeuteration of POPC is sufficient to induce phase separation. Perdeuteration of the DPPC chains pushes the system farther from the phase coexistence regime so that higher concentrations of melatonin (in this case 10 mol%) are required to promote phase separation.

## CONCLUSION

In this work we used ^2^H-NMR to study phase separation of POPC/DPPC/cholesterol model lipid membranes to understand the effects of melatonin on phospholipid bilayers. Melatonin induces *𝓁*_*o*_ : *𝓁*_*d*_ phase coexistence in the POPC/DPPC/cholesterol mixtures, in both POPC-*d*_31_ and DPPC-*d*_62_ containing samples, though the effect on POPC-*d*_31_ was observed 5 mol% of melatonin, which may imply that melatonin preferentially interacts with the unsaturated lipid (POPC). Based on our results and other relevant literature we suggest that a competition between melatonin and cholesterol in binding to lipid molecules results in the displacement of cholesterol by melatonin in the more disordered phase, which acts to increase the disorder in that phase, while driving more cholesterol into DPPC ordered phase and thus promoting the separation of *𝓁*_*o*_ : *𝓁*_*d*_ phases. POPC has a larger area per molecule than DPPC, therefore there is more free space in the headgroup region which allows more melatonin to incorporate into POPC enriched domains. In addition, the perdeuteration of lipid chains lowers the melting temperature of the corresponding lipid, which increases the difference between POPC-*d*_31_ and DPPC, and reduces the difference between DPPC-*d*_62_ and POPC. In this simple POPC/DPPC/cholesterol model membrane system, melatonin clearly acts to promote membrane fluid phase heterogeneity. Considering the importance of membrane nanodomains in amyloid toxicity reported in the earlier work by Drolle et al. (2, 29), this data will be important to guide future studies on melatonin’s protective role on a membrane level.

## AUTHOR CONTRIBUTIONS

All authors participated in designing the research, interpreting the results and preparing this manuscript. Nanqin Mei prepared samples with assistance from Morgan Robinson. Nanqin Mei and James H. Davis collected and analyzed all data.

## ACKNOWLEDGMENTS

The authors acknowledge funding from Natural Sciences and Engineering Research Council, grants to James H. Davis and Zoya Leonenko, and Ontario Graduate Scholarship to Morgan Robinson. We also thank the staff of the University of Guelph NMR Centre for their assistance.

## REFERENCES

1. Drolle, E., N. Kučerka, M. I. Hoopes, Y. Choi, J. Katsaras, M. Karttunen, and Z. Leonenko, 2013. Effect of melatonin and cholesterol on the structure of DOPC and DPPC membranes. Biochim. Biophys. Acta 1828:2247–2254.

2. Drolle, E., A. Negoda, K. Hammond, E. Pavlov, and Z. Leonenko, 2017. Changes in lipid membranes may trigger amyloid toxicity in Alzheimer’s disease. PloS one 12:e0182194.

3. Davis, J. H., J. J. Clair, and J. Juhasz, 2009. Phase equilibria in DOPC/DPPC-d62/cholesterol mixtures. Biophys. J. 96:521–539.

4. Martín, V., N. Fabelo, G. Santpere, B. Puig, R. Marín, I. Ferrer, and M. Díaz, 2010. Lipid alterations in lipid rafts from Alzheimer’s disease human brain cortex. Journal of Alzheimer’s Disease 19:489–502.

5. Chan, R. B., T. G. Oliveira, E. P. Cortes, L. S. Honig, K. E. Duff, S. A. Small, M. R. Wenk, G. Shui, and G. Di Paolo, 2012. Comparative lipidomic analysis of mouse and human brain with Alzheimer disease. J. Biol. Chem. 287:2678–2688.

6. Nicholson, A. M., and A. Ferreira, 2009. Increased membrane cholesterol might render mature hippocampal neurons more susceptible to beta-amyloid-induced calpain activation and tau toxicity. J. Neuroscience 29:4640–51.

7. Sepulveda, F. J., J. Parodi, R. W. Peoples, C. Opazo, and L. G. Aguayo, 2010. Synaptotoxicity of Alzheimer beta amyloid can be explained by its membrane perforating property. PLoS ONE 5:1–9.

8. Cecchi, C., and M. Stefani, 2013. The amyloid-cell membrane system. The interplay between the biophysical features of oligomers/fibrils and cell membrane defines amyloid toxicity. Biophys. Chem. 182:30–43.

9. Karasek, M., 2004. Melatonin, human aging, and age-related diseases. In Experimental Gerontology. volume 39, 1723–1729.

10. Uchida, K., N. Okamoto, K. Ohara, and Y. Morita, 1996. Daily rhythm of serum melatonin in patients with dementia of the degenerate type. Brain Research 717:154–159.

11. Liu, R. Y., J. N. Zhou, J. Van Heerikhuize, M. A. Hofman, and D. F. Swaab, 1999. Decreased melatonin levels in postmortem cerebrospinal fluid in relation to aging, Alzheimer’s disease, and apolipoprotein E-E 4*/*4*genotype*. J.Clin.84 : 323–327.

12. Cardinali, D., L. Brusco, C. Liberczuk, and A. Furio, 2002. The use of melatonin in Alzheimer’s disease. Neuroendocrinol. Lett. 23:20–23.

13. Singer, C., R. E. Tractenberg, J. Kaye, K. Schafer, A. Gamst, M. Grundman, R. Thomas, and L. J. Thal, 2003. A multicenter, placebo-controlled trial of melatonin for sleep disturbance in Alzheimer’s disease. Sleep 26:893–901.

14. Feng, Z., and J.-T. Zhang, 2004. Protective effect of melatonin on beta-amyloid-induced apoptosis in rat astroglioma C6 cells and its mechanism. Free Radic. Biol. Med. 37:1790–1801.

15. Pappolla, M. a., M. Sos, R. a. Omar, R. J. Bick, D. L. Hickson-Bick, R. J. Reiter, S. Efthimiopoulos, and N. K. Robakis, 1997. Melatonin prevents death of neuroblastoma cells exposed to the Alzheimer amyloid peptide. J. Neuroscience 17:1683–1690.

16. Chyan, Y. J., B. Poeggeler, R. A. Omar, D. G. Chain, B. Frangione, J. Ghiso, and M. A. Pappolla, 1999. Potent neuroprotective properties against the Alzheimer *β*-amyloid by an endogenous melatonin-related indole structure, indole-3-propionic acid. J. Biol. Chem. 274:21937–21942.

17. Ali, T., and M. O. Kim, 2015. Melatonin ameliorates amyloid beta-induced memory deficits, tau hyperphosphorylation and neurodegeneration via PI3/Akt/GSk3*β* pathway in the mouse hippocampus. J. Pineal Res. 59:47–59.

18. Olcese, J. M., C. Cao, T. Mori, M. B. Mamcarz, A. Maxwell, M. J. Runfeldt, L. Wang, C. Zhang, X. Lin, G. Zhang, and G. W. Arendash, 2009. Protection against cognitive deficits and markers of neurodegeneration by long-term oral administration of melatonin in a transgenic model of Alzheimer disease. J. Pineal Res. 47:82–96.

19. Pappolla, M., M. Simovich, T. Bryant-Thomas, Y. Chyan, B. Poeggeler, M. Dubocovich, R. Bick, G. Perry, F. Cruz-Sanchez, and M. Smith, 2002. The neuroprotective activities of melatonin against the Alzheimer beta-protein are not mediated by melatonin membrane receptors. J. Pineal Res. 32:135–142.

20. Dies, H., B. Cheung, J. Tang, and M. C. Rheinstädter, 2015. The organization of melatonin in lipid membranes. Biochim. Biophys. Acta 1848:1032–1040.

21. Ceraulo, L., S. Fanara, V. T. Liveri, A. Ruggirello, W. Panzeri, and A. Mele, 2008. Orientation and molecular contacts of melatonin confined into AOT and lecithin reversed micellar systems. Colloid. Surface A. 316:307–312.

22. Choi, Y., S. J. Attwood, M. I. Hoopes, E. Drolle, M. Karttunen, and Z. Leonenko, 2014. Melatonin directly interacts with cholesterol and alleviates cholesterol effects in dipalmitoylphosphatidylcholine monolayers. Soft Matter 10:206–213.

23. Bongiorno, D., L. Ceraulo, M. Ferrugia, F. Filizzola, A. Ruggirello, and V. T. Liveri, 2005. Localization and interactions of melatonin in dry cholesterol/lecithin mixed reversed micelles used as cell membrane models. J. Pineal Res. 38:292–298.

24. Stillwell, W., 2013. An introduction to biological membranes: from bilayers to rafts. Newnes.

25. Simons, K., and E. Ikonen, 1997. Functional rafts in cell membranes. Nature 387:569.

26. Nickels, J. D., S. Chatterjee, C. B. Stanley, S. Qian, X. Cheng, D. A. Myles, R. F. Standaert, J. G. Elkins, and J. Katsaras, 2017. The in vivo structure of biological membranes and evidence for lipid domains. PLoS Biol. 15:e2002214.

27. Davis, J. H., 1983. The description of membrane lipid conformation, order and dynamics by 2H-NMR. Biochim. Biophys. Acta 737:117–171.

28. Veatch, S. L., and S. L. Keller, 2003. Separation of Liquid Phases in Giant Vesicles of Ternary Mixtures of Phospholipids and Cholesterol. Biophys. J. 85:3074–3083.

29. Drolle, E., R. M. Gaikwad, and Z. Leonenko, 2012. Nanoscale electrostatic domains in cholesterol-laden lipid membranes create a target for amyloid binding. Biophysical journal 103:L27–L29.

30. Vist, M. R., and J. H. Davis, 1990. Phase equilibria of cholesterol/dipalmitoylphosphatidylcholine mixtures: deuterium nuclear magnetic resonance and differential scanning calorimetry. Biochemistry 29:451–464.

31. Davis, J. H., 1979. Deuterium magnetic resonance study of the gel and liquid crystalline phases of dipalmitoyl phosphatidylcholine. Biophys. J. 27:339–358.

32. Davis, J. H., K. Jeffrey, M. Bloom, M. Valic, and T. Higgs, 1976. Quadrupolar echo deuteron magnetic resonance spectroscopy in ordered hydrocarbon chains. Chem. Phys. Letters 42:390–394.

33. van Gorkom, L. C. M., J. M. Hook, M. B. Logan, J. V. Hanna, and R. E. Wasylishen, 2013. Solid-state lead-207 NMR of lead(II) nitrate: Localized heating effects at high magic angle spinning speeds. Magn. Reson. Chem. 33:791–795.

34. Beckmann, P. A., and C. Dybowski, 2000. A thermometer for nonspinning solid-state NMR spectroscopy. J. Magn. Reson. 146:379–380.

35. Schmidt, M. L., L. Ziani, M. Boudreau, and J. H. Davis, 2009. Phase equilibria in DOPC/DPPC: Conversion from gel to subgel in two component mixtures. J. Chem. Phys. 131:175103.

36. Marsh, D., 2013. Handbook of lipid bilayers. CRC press.

37. Tardieu, A., V. Luzzati, and F. Reman, 1973. Structure and polymorphism of the hydrocarbon chains of lipids: a study of lecithin-water phases. J. Mol. Biol. 75:711–733.

38. Davis, J. H., and M. L. Schmidt, 2019. Cholesterol in Model Membranes, Walter de Gruyter GmbH, 325–364. https://www.degruyter.com/viewbooktoc/product/488626.

39. Zhao, J., J. Wu, H. Shao, F. Kong, N. Jain Gunt, and G. Feigenson, 2007. Phase studies of model biomembranes: Macroscopic coexistence of Lα +. L*β*b, with light-induced coexistence of Lα + Lo phases. Biochim Biophys. Acta 1768:2777–2786.

40. Konlyakhina, T. M., S. L. Goh, J. Amazon, F. A. Heberle, J. Wu, and G. W. Feigenson, 2011. Control of a nanoscopic-to-macroscopic transition: Modulated phases in four-component DSPC/DOPC/POPC/Chol giant unilamellar vesicles. Biophys. J. 101:L08–L20.

41. Goh, S. L., J. J. Amazon, and G. W. Feigenson, 2013. Toward a better raft model: Modulated phases in the four-component bilayer, DSPC/DOPC/POPC/CHOL. Biophys. J. 104:853–862.

42. Heberle, F. A., R. S. Petruzielo, J. Pan, P. Drazba, N. Kucerka, R. F. Standaert, G. W. Feignson, and J. Katsaras, 2013. Bilayer thickness mismatch controls domain size in model membranes. J. Am. Chem. Soc. 135:6853–6859.

43. Kučerka, N., M.-P. Nieh, and J. Katsaras, 2011. Fluid phase lipid areas and bilayer thicknesses of commonly used phosphatidylcholines as a function of temperature. Biochimica et Biophysica Acta (BBA)-Biomembranes 1808:2761–2771.

