## Supplemental File for "Melatonin alters fluid phase co-existence in POPC/DPPC/cholesterol membranes"

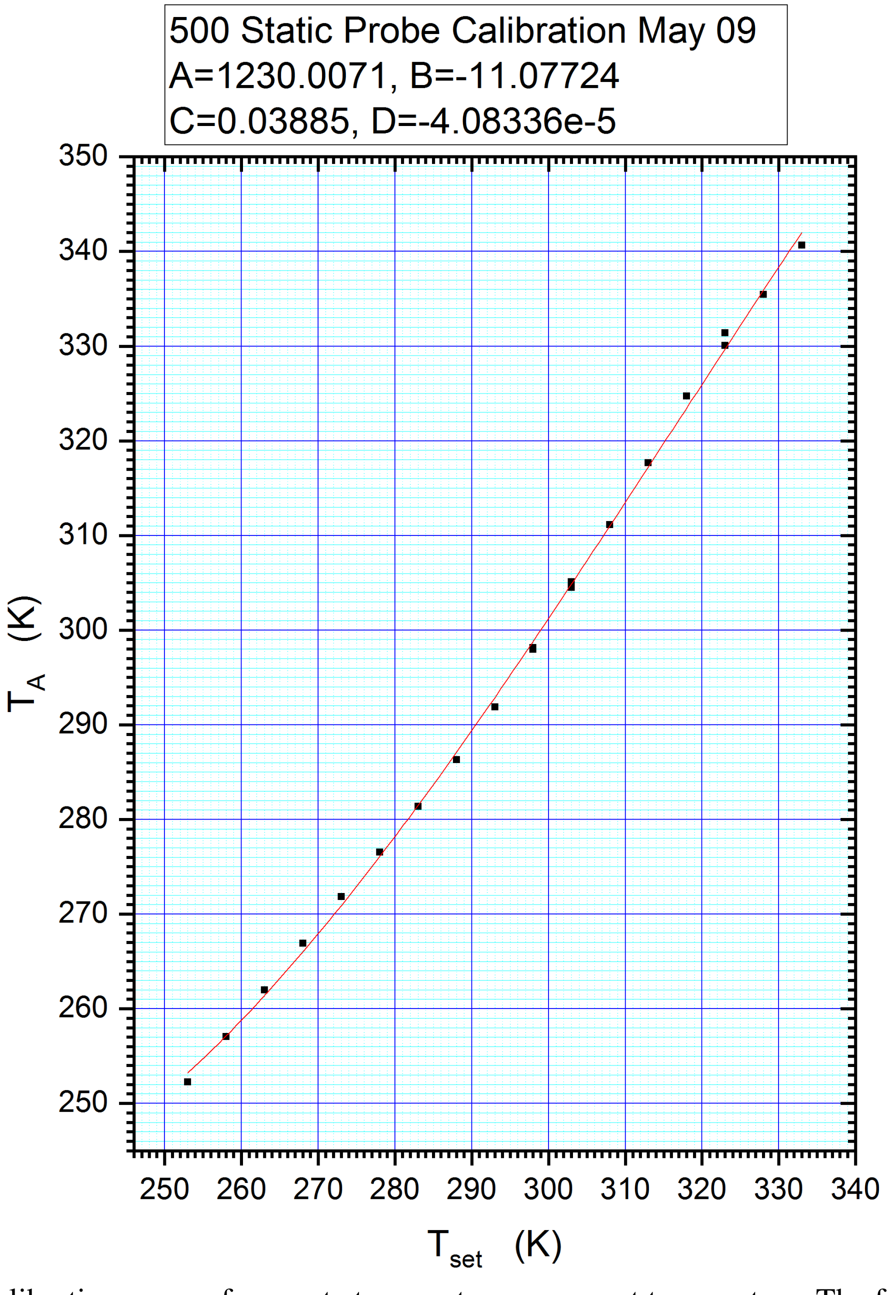


Figure 1. Calibration curve of accurate temperature versus set temperature. The function of the fitted curve is expressed as: $T_A = A + B * T_{set} + C * {T_{set}}^2 + D * {T_{set}}^3$.
